# Plant phase extraction (PPE): A novel method for enhanced discovery of RNA-binding proteome in plants

**DOI:** 10.1101/2022.06.02.494555

**Authors:** Yong Zhang, Ye Xu, Todd H. Skaggs, Jorge F.S. Ferreira, Xuemei Chen, Devinder Sandhu

## Abstract

RNA-binding proteins (RBPs) are versatile effectors in posttranscriptional gene regulation. Systematic profiling of RBPs in plants has been limited to proteins interacting with polyadenylated (poly(A)) RNAs due to the lack of an efficient method of isolating RBPs associated with non-poly(A) RNAs. Here we reported the establishment and application of plant phase extraction (PPE) as a novel method to comprehensively discover the RNA-binding proteome in Arabidopsis leaf tissues, leading to the isolation of 1,169 RBPs, of which 673 corroborate with previously reported RBPs and 496 are novel RBPs. PPE showed unmatched ability in capturing 374 diverse RNA-binding domains (RBDs), while 44% of the RBPs lack recognized RBDs. PPE recovered far more ribosomal and tRNA-related RBPs and metabolic enzymes with moonlighting RBP functions than any other individual report from plants, signifying its advantage in unbiasedly retrieving non-poly(A) and unconventional RBPs. We proposed that intrinsically disordered regions (IDRs) contributed to the non-classical binding of some novel RBPs, and provided the first evidence that enzymatic domains from metabolic enzymes have additional roles in RNA-binding. Taken together, our findings demonstrated that PPE is an effective approach to identifying a wide range of RBPs in complex plant tissues and may have broad biological implications.

## Introduction

RNA-binding proteins (RBPs) form dynamic ribonucleoprotein (RNP) complexes with RNA and play vital roles in posttranscriptional gene regulation. Virtually every aspect of RNA metabolism, including maturation (capping, splicing, and polyadenylation), editing, modification, subcellular localization, translation, and degradation, is exquisitely modulated by a myriad of RBPs that coat RNA transcripts in a general or spatiotemporal manner (Glisovic et al., 2008; Singh et al., 2015). In humans, defective RBPs cause cancer, metabolic disorders, and neurodegenerative diseases (Verkerk et al., 1991; Lefebvre et al., 1995; Bechara et al., 2013; Patel et al., 2015; Wang et al., 2019; Gebauer et al., 2021; Louis et al., 2021). In plants, RBPs are crucial regulators implicated in ovule development, floral transition, circadian rhythm, stress, and immune responses (Heintzen et al., 1997; Macknight et al., 1997; Schomburg et al., 2001; Kim et al., 2005; Zhang et al., 2005; Deleris et al., 2006; Zhang et al., 2011; Bush et al., 2015; Zhang et al., 2015; Albaqami et al., 2019; Marondedze et al., 2019a). Therefore, investigating interplays between RBPs and RNA has undoubtedly shaped and deepened our understanding of how RBPs govern developmental processes and environmental responses.

Typical RBP-RNA interactions are mediated by RNA-binding domains (RBDs), which recognize either specific sequence elements or secondary/tertiary structures of their RNA partners. The most prevalent RBDs are conserved in eukaryotic cells, including the RNA recognition motif (RRM), the K-homology (KH) domain, the DEAD-box helicase domain, double-stranded RNA-binding motif (DSRM), zinc-finger domain, and other less abundant domains (Silverman et al., 2013; Gerstberger et al., 2014). It is worth noting that plant RBPs are more diversified than their mammalian counterparts. For instance, about 50% of RRM-type plant RBPs have no ortholog in metazoans (Lorkovic and Barta, 2002). Also, far more predictable plant pentatricopeptide repeats (PPR)-type RBPs, which regulate posttranscriptional events in plastids and mitochondria (Cheng et al., 2016), suggest that these RBPs are more likely involved in plant-specific biological processes.

Classical RBDs have enabled computational prediction of RBPs based on sequence homology across species. However, mounting evidence demonstrated that proteins without known RBDs are capable of binding RNAs as well, including those metabolic enzymes such as thymidylate synthase, aconitase, and glyceraldehyde 3-phosphate dehydrogenase (GAPDH) with moonlighting RBP functions that are distinct from their basic role in metabolism (Chu et al., 1991; Hentze and Argos, 1991; Chang et al., 2013). The recurrent identification of unconventional RBPs strongly indicated that the number of RBPs has been underestimated, and the way RBPs interact with RNAs is more sophisticated than predicted. Indeed, the human RBP repertoire has immensely expanded, with about half of the RBPs lacking canonical RBDs revealed by RNA interactome capture (RIC) in human cell lines (Baltz et al., 2012; Castello et al., 2012). RIC is a method based on UV-crosslinking and oligo(dT) affinity purification to systematically profile RNA-binding proteome (RBPome) bound to polyadenylated (poly(A)) RNAs (Baltz et al., 2012; Castello et al., 2012). This powerful approach has since been applied to various cell types and organisms, leading to the discovery of 1,914 RBPs in human, 1,393 in mouse, 1,273 in yeast, 777 in Drosophila, and 594 in nematode (Kwon et al., 2013; Beckmann et al., 2015; Matia-Gonzalez et al., 2015; Liao et al., 2016; Liepelt et al., 2016; Sysoev et al., 2016; Despic et al., 2017; Hentze et al., 2018). RIC has later been adapted and improved in Arabidopsis using etiolated seedlings, mesophyll protoplasts, root cell cultures, and leaves (Marondedze et al., 2016; Reichel et al., 2016; Zhang et al., 2016; Marondedze et al., 2019b; Bach-Pages et al., 2020). When combined, a staggering number of RNA-related proteins were identified (2,701), with 836 proteins registered as RBPs and 1,865 proteins as candidate RBPs in Arabidopsis (Marondedze, 2020). However, the overlap between these datasets is considerably low, as all four Arabidopsis RBPomes share only 25 RBPs (Bach-Pages et al., 2020).

RIC has been extensively explored in eukaryotes throughout the past decade; however, very few of these newly discovered proteins were experimentally validated as *bona fide* RBPs. More importantly, this approach is heavily biased toward RBPs that bind poly(A) RNAs. Given that mRNA only represents less than 5% of the population of total RNA in eukaryotic cells, the full spectrum of RBPs might still be far from complete due to the incompetent recovery of RBPs that specifically interact with the vast majority of non-poly(A) RNAs. To circumvent the inherent limitation of RIC, two methods, namely RICK (capture of the newly transcribed RNA interactome using click chemistry) and CARIC (click chemistry-assisted RNA interactome capture), based on metabolic labeling of cells with nucleotide analogs followed by click chemistry and streptavidin affinity enrichment, were used to characterize RBPs attached to both coding and non-coding RNAs (Bao et al., 2018; Huang et al., 2018). Although these approaches addressed the drawback of RIC by recovering RBPs that potentially bind to non-ploy(A) RNAs, they rely on efficient *in vivo* labeling of RNA and can introduce bias caused by transcription-dependent nucleotide incorporation. Additionally, the application of these methods to complex tissues is hindered due to the limited uptake of nucleotide analogs.

New strategies using phase separation/extraction, including XRNAX (protein-crosslinked RNA extraction), OOPS (orthogonal organic phase separation), and PTex (phenol toluol extraction), have been developed to unbiasedly identify RBPs in human cell lines, mouse brain, and bacteria (Queiroz et al., 2019; Trendel et al., 2019; Urdaneta et al., 2019). Crosslinked RBP-RNA adducts can be separated from free RNA and proteins solely based on their physicochemical properties, using acid guanidinium thiocyanate-phenol (commercially available as Trizol reagent)-chloroform phase extraction. The crosslinked RBP-RNA adducts migrate into the insoluble interphase, free RNA moves to the aqueous phase, and proteins remain in the organic phase, thereby overcoming the challenges mentioned in the foregoing approaches. RBPs, regardless of their RNA type, including many of those discovered by RIC and hundreds of new ones that bind non-poly(A) RNAs were captured using these approaches (Queiroz et al., 2019; Trendel et al., 2019; Urdaneta et al., 2019).

Phase extraction, such as OOPS, was also attempted in Arabidopsis (Liu et al., 2020). Unfortunately, the result fell short of expectations and was overshadowed by previous RIC studies using similar plant materials (Marondedze et al., 2016; Bach-Pages et al., 2020). Among the 468 proteins identified by OOPS, only 244 were considered “RBP”, and the rest remained debatable. Furthermore, only a small fraction of these putative RBPs was shared by other studies, raising the question if plant RBPs can be effectively separated by phase extraction. Inspired by the successful application of phase extraction in human cell lines and bacteria, here we developed plant phase extraction (PPE), customized for plant tissue samples. Using PPE, we identified 1,169 RBPs, including traditional RBPs participating in various aspects of RNA metabolism and a plethora of metabolic enzymes with moonlighting RBP functions. Remarkably, we discovered 496 novel RBPs, many of which are engaged in non-coding RNA binding. We showed that intrinsically disordered regions (IDRs) widely occurred in the PPE-RNA-binding proteome (PPE-RBPome) and may be partially responsible for non-canonical RNA-binding. Moreover, putative novel RBDs from non-classical RBPs were revealed, and a selected set was validated as novel RBDs for the first time.

## Results

### Development of plant phase extraction (PPE) in Arabidopsis

Since plant tissues are more complex than human cell lines due to cell walls, plastids, and numerous secondary metabolites, we speculated that these could have caused the poor outcome using OOPS in Arabidopsis. We recapitulated the OOPS procedures as described (Liu et al., 2020) with three cycles of biphasic separation using Trizol to directly lyse leaf cells from fine-powdered samples ground in liquid nitrogen). In striking contrast to the thin layer of sticky white interphase, predominantly composed of RBP-RNA adducts seen in human cells, phase extraction of Arabidopsis leaves resulted in a thicker and solid interphase layer (Supplemental Figure S1). The interphase layer consisted of cell debris, starch, secondary metabolites, RBP-RNA adducts, and other molecules tightly associated with cell debris, making it impractical to separate the embedded RBP-RNA adducts from the contaminants without compromising their purity and quantity. We, therefore, tailored the XRNAX method for Arabidopsis with several critical modifications and designated this strategy as plant phase extraction (PPE). Firstly, to counteract the negative effects of UV-absorbing pigments from leaves, three cycles of UV-crosslinking were conducted at 200 mJ/cm^2^ (254 nm wavelength), irradiating the adaxial side twice and the abaxial side once (Zhang et al., 2015; Bach-Pages et al., 2020). Secondly, a lysis buffer containing polyvinylpyrrolidone 40 was added to plant tissue ground in liquid nitrogen to remove secondary metabolites, which would interfere with mass spectrometry analysis. Then, centrifugation was performed before adding Trizol and chloroform to the supernatant to remove the cell debris and other insoluble substances prior to the capture of RBP-RNA adducts. Thirdly, since the interphase was not as sticky as that originated from human cells, it was difficult to wash away the residual aqueous and organic phases without disrupting the integrity of the interphase. To avoid unnecessary loss of RBPs caused by washing, we replaced the washing step used in XRNAX with two rounds of phase extraction. Lastly, after DNase digestion and before isopropanol precipitation of the RBP-RNA adducts, two more rounds of phase extraction were added to remove DNase and carry-overs from previous steps as much as possible (Figure 1A).

**Fig. 1.**
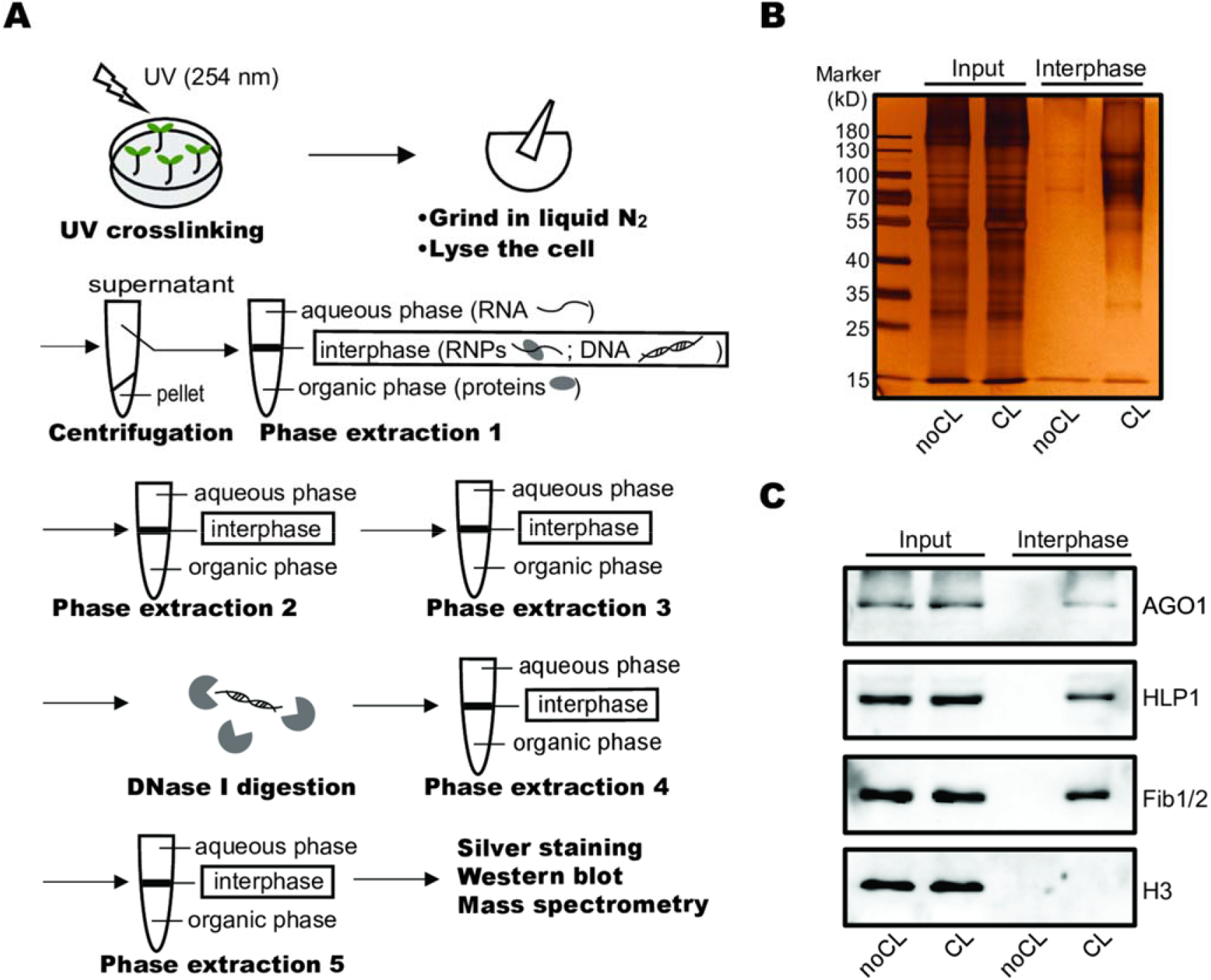
Plant Phase Extraction (PPE) in Arabidopsis. **(A)** Schematic representation of PPE. **(B)** Silver staining of the SDS-PAGE gel with interphase proteins acquired by PPE alongside with diluted input (20-fold dilution) from non-crosslinked (noCL) and crosslinked (CL) samples. **(C)** Western blot showing enriched RBPs (AGO1, HLP1 and Fib1/2), but not histone H3 in the CL samples.

To evaluate the efficiency of PPE, we prepared interphases from non-crosslinked (noCL) and crosslinked (CL) samples (12-day-old leaves of Arabidopsis). A thin insoluble interphase layer similar to the one shown in human cells was recovered from each sample, and the difference in the interphase layer between noCL and CL samples can be easily observed (Supplemental Figure S1). Proteins from these interphase layers were separated by SDS-PAGE and visualized by silver staining (Figure 1B). Since certain groups of proteins, such as glycosylated proteins, share the physicochemical properties of RNA-protein adducts, they are present in the interphase layers from both CL and noCL samples (Queiroz et al., 2019). However, compared to noCL sample, more proteins were enriched in the CL sample with the same amount of starting material (Figure 1B). Western blot using antibodies against several known RBPs, AGO1, HLP1, and Fib1/2 (Song et al., 2003; Rakitina et al., 2011; Zhang et al., 2015), further confirmed the presence of RBPs in the CL sample but not in the noCL control (Figure 1C). As an abundant non-RBP, histone H3 was not detected in the CL sample. These results demonstrated PPE to be an efficient way to purify RBPs from plant tissues.

### Comprehensive identification of the Arabidopsis leaf RBPome by PPE

Having established PPE to enrich RBPs in Arabidopsis, we sought to identify the RNA-binding proteome (RBPome) in Arabidopsis leaf tissue using mass spectrometry. The reproducibility among three replicates was demonstrated by scatter plots comparing the protein enrichment in CL samples over noCL samples (log_2_ fold change [CL/noCL], Figure 2A). Proteins present in at least two out of the three replicates with a fold change [CL/noCL] ≥ 2 at an adjusted *p*-value (FDR) < 5% were defined as RBPs, and those with a fold change [CL/noCL] ≥ 1.5 at an FDR < 20% were considered as candidate RBPs (Backlund et al., 2020) (Figure 2A). We identified 1,169 RNA-associated proteins, of which 1,118 proteins were classified as RBPs and 51 as candidate RBPs (Supplemental Data Set 1). For simplicity, both RBPs and candidate RBPs were grouped for the following analyses. According to current gene ontology (GO) annotations, 49% of these proteins are linked to RNA biology; the rest, 51%, have no previously assigned RNA-related functions (Figure 2B). GO enrichment analysis showed an over-representation of RNA-binding, ribosomal-related, and tRNA-related terms, such as tRNA ligase, tRNA editing, and tRNA binding activities (Figure 2C). We noticed that “amino acid activation” and “ncRNA metabolic process” topped the most enriched GO terms in “biological process”, highlighting the efficiency of PPE in recovering aminoacyl-tRNA ligase and other non-coding RNA-binding proteins (Supplemental Figure S2). Some of the most enriched terms in the “RNA binding” subcategory, such as “rRNA binding”, “tRNA binding”, “snoRNA binding,” and “7S RNA”, further denoted that RBPs bound to non-coding RNAs were successfully discovered using PPE (Figure 2D). RBPs were localized to various cellular compartments (Supplemental Figure S3A), with an over-representation of GO terms in the chloroplast, cytoplasm (ribosome, stress granule, peroxisome, etc.), and nucleolus, in line with their fundamental roles in RNA processing, transport, and translation (Supplemental Figure S3B).

**Fig. 2.**
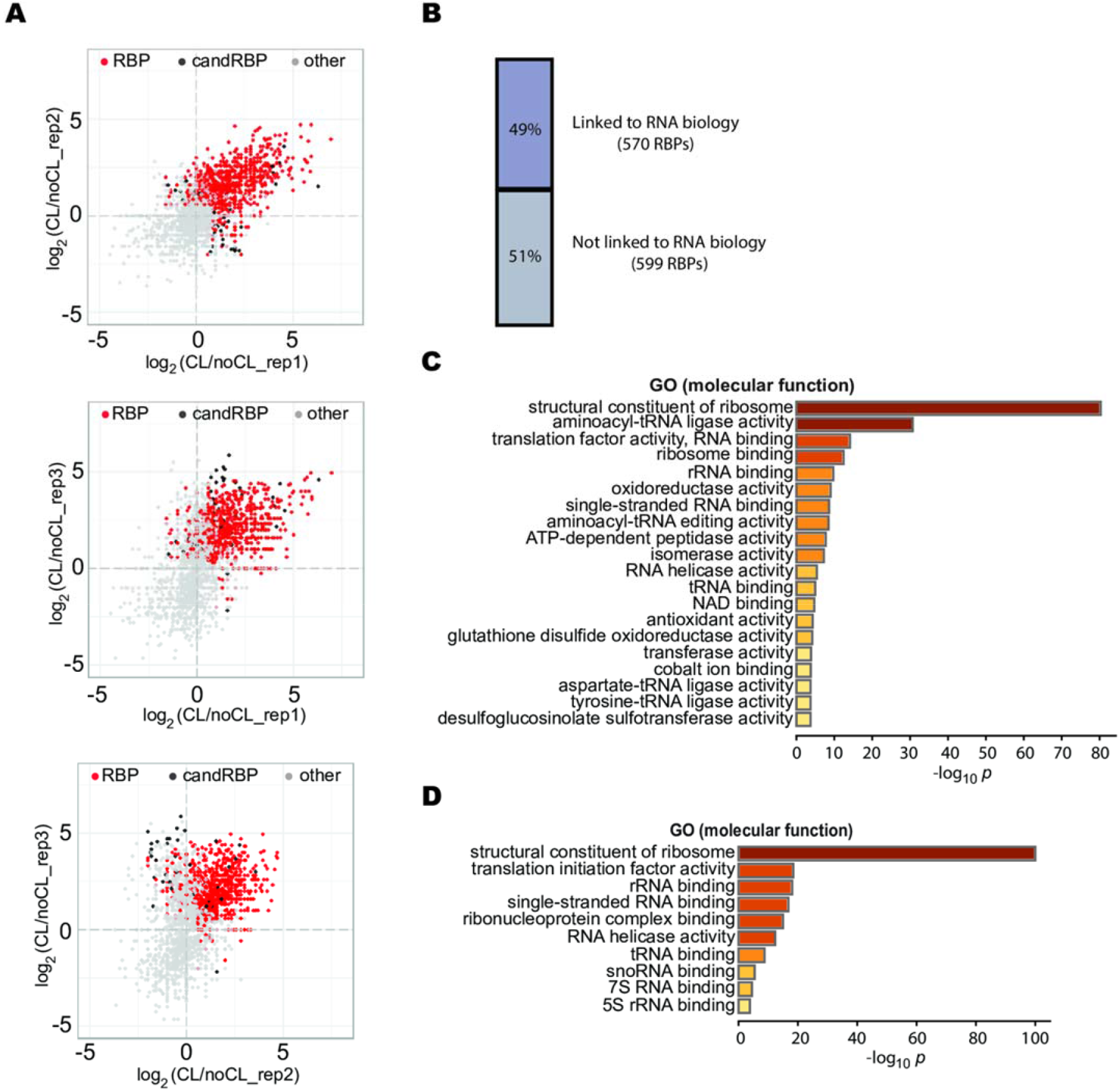
Identification of the RBP proteome using PPE. **(A)** Scatter plots showing the reproducibility between biological replicates by comparing the log_2_ fold change [CL+1/noCL+1]. Statistically enriched RBPs and candidate RBPs (candRBP) are indicated in red and black, respectively. Gray dots represent non-RBPs. **(B)** Proportions of the RBPs that are linked to RNA biology (52%) and not linked to RNA biology (48%) based on gene ontology (GO) annotation. Numbers of each category are also shown. (**C)** Top twenty enriched molecular function GO terms of the proteome identified using PPE. (**D)** Over-represented non-coding RNA binding terms in the RNA binding subcategory.

### RBPs identified by PPE display a broad array of RNA-binding domains

To evaluate potential RBDs possessed by the 1,169 RBPs, we first integrated published RBDs with experimental evidence (Castello et al., 2012; Castello et al., 2016) and retrieved a total of 1,152 Pfam accessions covering both classical and non-classical RBDs as the reference RBD database (Supplemental Data Set 2). About 32% of the RBDs in the database (374) can be found in 56% of the PPE-RBPs (650 RBPs), leaving the remaining 519 RBPs without recognizable RBDs in the current database (Figure 3A). Compared to RIC and OOPS, PPE identified more RBDs (Figure 3B). For RBPs with known RBDs, GO enrichment analysis showed RNA binding activities towards all major types of RNAs, such as “single-stranded mRNA binding”, “rRNA binding”, “tRNA binding”, and “snoRNA binding”. The most over-represented “molecular function” terms included “structural constituent of ribosome”, “aminoacyl-tRNA ligase activity”, and “translation factor activity”, implicating a major role of these RBPs in the translation process (Supplemental Figure S4A). For RBPs without known RBDs, GO terms were overwhelmingly enriched in metabolic activities, such as oxidoreductase, transferase and isomerase, and other non-RNA binding activities (Supplemental Figure S4B).

**Fig. 3.**
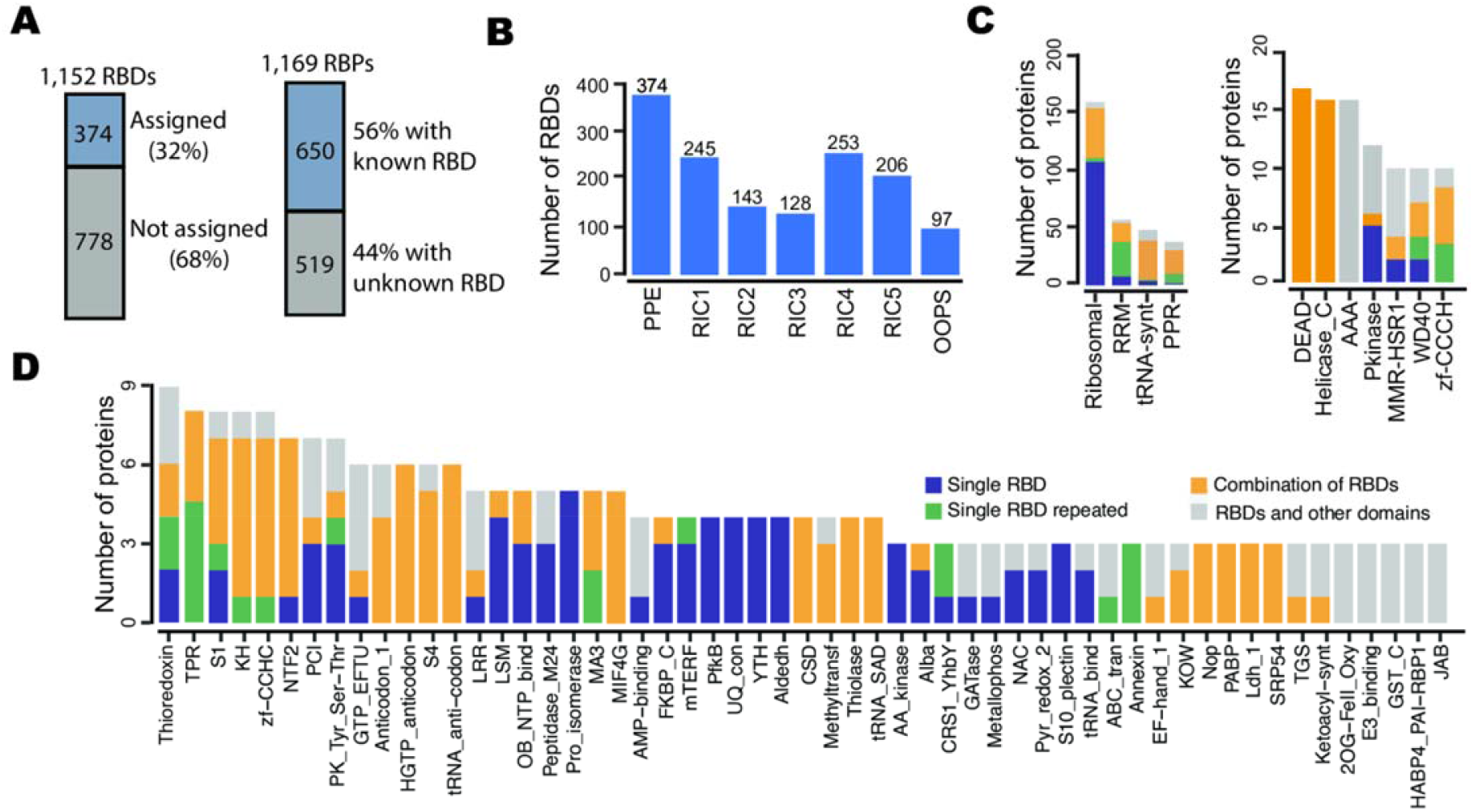
PPE identified a wide range of RNA-binding domains (RBDs). **(A)** Numbers and percentages of known RBDs that can be assigned to RBPs are shown. In the database with 1,152 RBDs, 374 (32%) can be assigned to 650 RBPs. Numbers and proportions of RBPs with known (650/1,169, 56%) and unknown RBDs (519/1,169, 44%) are shown. **(B)** PPE identified more RBDs than RIC and OOPS. RIC1 (5/6-week-old leaves); RIC2 (4-week-old leaves); RIC3 (4/5-week-old mesophyll protoplast); RIC4 (4-day-old etiolated seedlings); RIC5 (root suspension cell); OOPS (4-week-old whole plant). (**C)** Most abundant RBDs (number of proteins ≥ 10). (**D)** Abundant RBDs (3∼9 proteins). For **(C)** and **(D)**, RBPs with a single RBD, with more than one same type of RBDs, with two or more different types of RBDs, or have a combination of RBD(s) and non-RBD(s) are depicted in blue, green, yellow and gray, respectively.

As a common feature, RBPs can harbor one or multiple RBDs (of either the same type or a combination of different types of RBDs) with or without the occurrence of other non-RBDs, to increase binding affinity and specificity (Lunde et al., 2007). Among the RBD classes, only ten have ≥10 protein members (Figure 3B); 55 have 3∼9 protein members (Figure 3C); the majority of them have only one or two members (Supplemental Figure S5). Classical RBDs, such as RRM, DEAD, KH, zf-CCCH, and cold shock domain (CSD), were identified (Figure 3B and3C). The most represented RBDs were the 185 ribosomal-related RBDs that originated from 160 proteins, followed by 116 RRMs from 57 RBPs, 48 tRNA-synt domains from 37 proteins, and 203 PPRs from 31 proteins (Figure 3B). Known non-classical RBDs, such as Helicase_C, AAA, MMR_HSR1, WD40, and TPR, were all captured. Notably, PPE uncovered nine thioredoxin-containing proteins. Although the N-terminus of human thioredoxin (TXN) was shown to interact directly with RNA (Castello et al., 2016), the thioredoxin domain in plants has not been registered as RBD yet. Domain analysis of 16 thioredoxin-containing proteins identified by PPE and RIC clearly showed that thioredoxin is the only domain shared by all the proteins, suggesting that thioredoxin is a novel RBD in plants (Figure 3C and Supplemental Figure S6). Similarly, we characterized domains, such as PKinase and PK_Tyr_Ser_Thr, from protein kinases that were shown to bind RNA in humans (Castello et al., 2016); therefore, they represent novel plant RBDs as well (Figures 3B and 3C). In addition, RBDs involved in non-coding RNA binding were identified, which include the Fibrillarin and Gar1 domains for snoRNAs, tRNA_binding domain for tRNAs, PAZ and Piwi domains for microRNAs and small interfering RNAs, and SRP54, the conserved subunit of the signal recognition particle (SRP) which interacts with 7S RNA (see Figure 3B, 3C and Supplemental Data Set 2 for a full list of RBDs identified by PPE).

### PPE identified novel RBPs and RBDs

Since earlier studies in Arabidopsis used different materials, UV crosslinking conditions, and statistical criteria to define RBPs, only 25 RBPs were exhibited in all datasets (Bach-Pages et al., 2020). Therefore we decided to compile all the published Arabidopsis RBPs/candidate RBPs acquired by RIC (Marondedze, 2020) and OOPS (Liu et al., 2020), irrespective of the differences in cell or tissue type, developmental stage, or treatment applied, to compare with RBPs discovered by PPE. Of the 1,169 RBPs identified by PPE, 673 were described previously as “RBP” or “candidate RBP” (Figure 4A and Supplementary Data Set 3). Meanwhile, 330 proteins formerly marked as “candidate RBP” were confirmed to be RBPs by our study (Supplementary Data Set 3). Strikingly, 496 characterized proteins out of 1,169 have never been reported as “RBP” in Arabidopsis, hence representing novel RBPs.

**Fig. 4.**
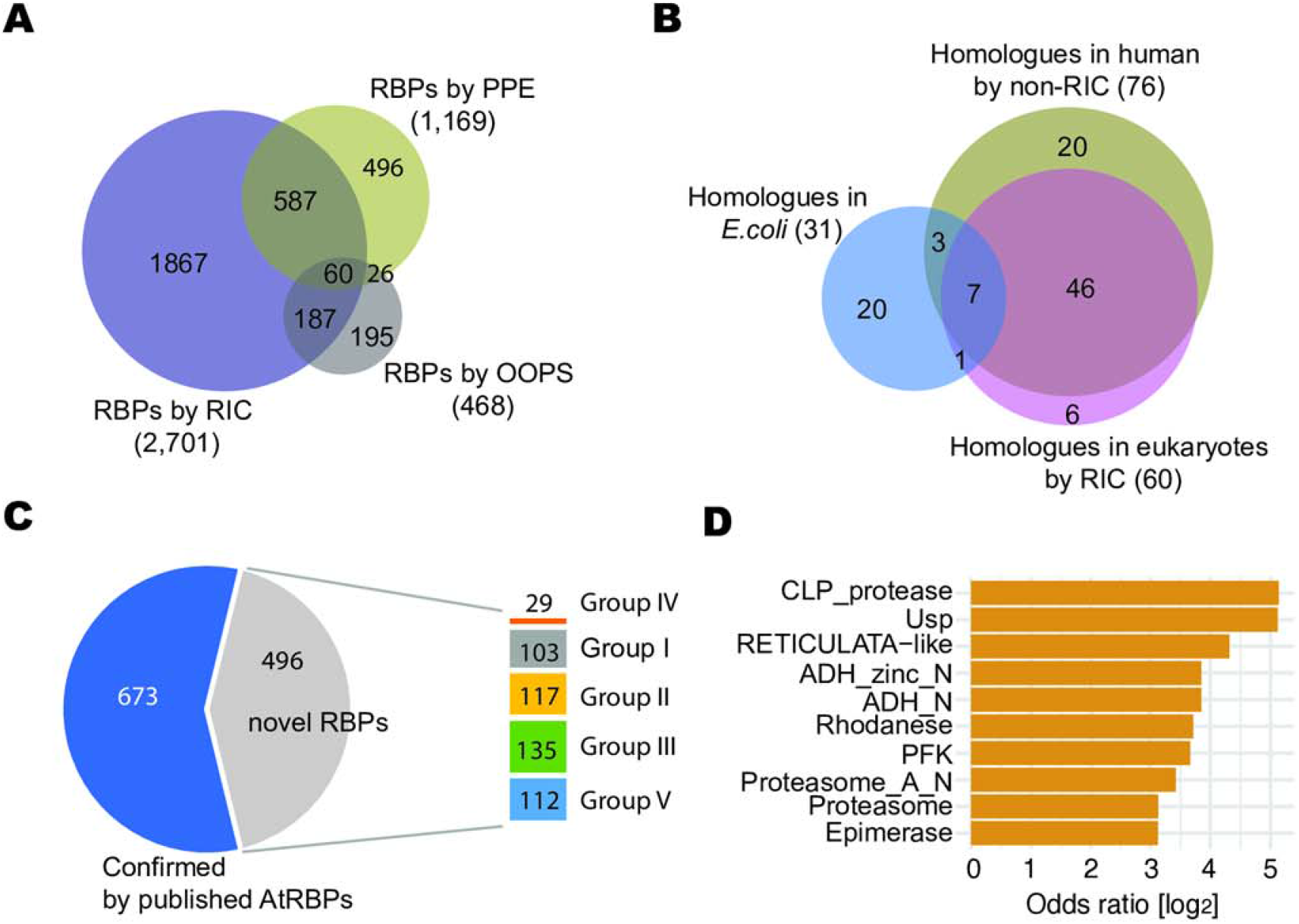
Novel RBPs identified by PPE. **(A)** Comparison of RBPs identified by PPE and other methods (RIC and OOPS) in Arabidopsis. PPE confirmed 673 RBPs classified by RIC and OOPS, and presented 496 novel RBPs. (**B)** A total of 103 proteins out of the 496 newly identified RBPs have homologues. Total numbers of homologues in *E. coli* (31), eukaryotes (humans, mice, Drosophila, nematodes, and yeast) by RIC method (60), in humans using non-RIC method (76) and overlapping numbers were shown. (**C)** Detailed categorization of RBPs identified by PPE. Aside from the 673 RBPs overlapping with reported ones, the 496 novel RBPs were assigned into five groups: I. 103 RBPs with homologues in other organisms; II. 117 RBPs with RBDs; III. 135 RBPs with other members in the same protein family recognized as RBPs; IV. 29 RBPs with at least two family members identified; and V. other types of RBPs. **(D)** Over-represented domains as putative novel RBDs (Fisher’s exact test, *p* <0.05).

To gain more evidence supporting these 496 proteins to be actual RBPs, we first sought to investigate whether their homologs have been already defined as “RBPs” in other eukaryotes, assuming that the RNA-binding ability of these RBPs is conserved across species. We searched RBPs obtained by RIC in humans, mice, drosophila, nematodes, and yeast (Hentze et al., 2018) and discovered that 60 out of the 496 proteins have at least one counterpart in these organisms (Figure 4B and Supplementary Data Set 3). Next, we extended the search to human RBPs identified by non-RIC approaches, including RICK (Bao et al., 2018), CARIC (Huang et al., 2018), and phase extraction methods [XRNAX (Trendel et al., 2019), OOPS (Queiroz et al., 2019), and PTex (Urdaneta et al., 2019)]. In this search, 76 out of the 496 proteins, including 53 captured by RIC, were shown to have human equivalents (Figure 4B and Supplementary Data Set 3). Lastly, we compared the remaining proteins with the *Escherichia coli* RBPome (Queiroz et al., 2019), as plants have their unique, bacteria-like chloroplast genome in addition to the nuclear and mitochondrial genomes. We found that 31 RBPs in this group, most localized in chloroplasts, were conserved in *E. coli* (Figure 4B and Supplementary Data Set 3). Collectively, our analyses revealed 103 non-redundant proteins with homology to RBPs in other organisms (designated as Group I).

In the remaining 393 proteins, 117 have defined RBDs according to the RBD reference database (designated as Group II; Figure 4C and Supplemental Data Set 4), thus representing high confident RBPs as well. Since proteins in the same family generally have similar structures and functions, we then focused on the rest 276 proteins to see whether they have family members classified as RBPs by RIC. If so, they are likely RBPs. In total, 135 proteins representing 75 protein families have members registered as “RBP” or “candidate RBP” (designated as Group III, Figure 4C and Supplemental Data Set 4).

We discovered additional 29 proteins from twelve protein families with at least two members in each family (Group IV). Although these proteins have no corresponding family members reported as RBPs in Arabidopsis so far, they might harbor certain sequences or structures that can serve as binding docks for RNA rather than being random contaminants because members from the same protein family could be repeatedly captured (Figure 4C and Supplemental Data Set 4). Domains shared within members of each protein family in Groups III and IV were summarized as putative novel RBDs (Supplemental Data Set 6). Enrichment analysis of the 432 Pfam accessions from the 496 novel RBPs by Fisher’s exact test further showed over-represented domains (*p* < 0.05).

Proteins in Group V do not directly fit the above-described criteria to be an RBP; however, these proteins contain domains similar to recognized RBDs. For instance, FLU (encoded by AT3G14110) harbors the TPR-like domain, and two thioredoxin protein family members (encoded by AT5G65840 and AT5G03880) contain the thioredoxin-like_sf and GST_N_3 domains, respectively (Supplemental Data Set 4). Although these domains have not been registered as RBDs, the similarity in sequence/structure compared to TPR and thioredoxin domains suggested that they are putative RBDs. Proteins involved in RNA biology, such as RNA degradation (SAL1/FRY1 and AT5G24340), tRNA modification (AT3G11800 and AT1G44835), and translation initiation (AT3G43540) were also discovered in this group, implying a role in RNA-binding (Supplemental Data Set 4).

### Intrinsically Disordered Regions in the PPE-RBPome contribute to RNA-binding

Intrinsically disordered regions (IDRs) are unstructured polypeptide segments and play critical roles in RBPs (Calabretta and Richard, 2015). Nearly half of the RNA-binding sites in humans occurred in IDRs (Castello et al., 2016). Therefore we speculated that IDRs residing in the PPE-RBPome might contribute, in part, to the non-canonical binding. We first retrieved 16,477 predicted IDRs in the Arabidopsis proteome from MobiDB, a database of protein disorder and mobility annotations (Piovesan et al., 2021), then located 634 IDRs to 475 RBPs. These include 325 RBPs confirmed by both PPE and RIC, and 150 novel ones, representing approximately 30% of the total novel RBPs (496) identified in the PPE-RBPome (Figure 5A and Supplemental Data Set 5). Specifically, we focused on groups IV and V proteins that lacked strong evidence for their justification to be “RBP” and uncovered 41 proteins with IDRs (7 in group IV and 34 in group V, Figure 5A). IDRs are widely immersed in protein-protein, protein-DNA, and protein-RNA interactions. We used flDPnn, by far one of the most accurate tools for coupling prediction of IDRs and their putative functions (Hu et al., 2021), to pinpoint whether these IDRs are engaged in RNA-binding rather than in protein-protein interaction or DNA binding. Most of the IDRs from both groups (six out the seven RBPs in group IV and 31 out of the 34 RBPs in group V with IDRs) have the potential to interact with RNA, as indicated by the propensity on the scale of 0 to 1 (Supplemental Data Set 7). For example, an uncharacterized protein encoded by AT4G16060 from group IV harbors an N-terminal (from amino acid 26 to 48) as well as a C-terminal (amino acid 271 to 289) disordered regions, both of which were predicted by flDPnn to confer RNA-binding ability (Figure 5B). Notably, we showed a stress-response protein encoded by AT3G59800 in group V is predicted to be in a completely unstructured form using AlphaFold (Figure 5C). The whole protein is disordered and displays a high propensity for RNA-binding, particularly at its N-terminus and C-terminus (Figure 5D). Taken together, our analyses implicated a role for IDRs in mediating interactions between RNA and the novel RBPs identified by PPE.

**Fig. 5.**
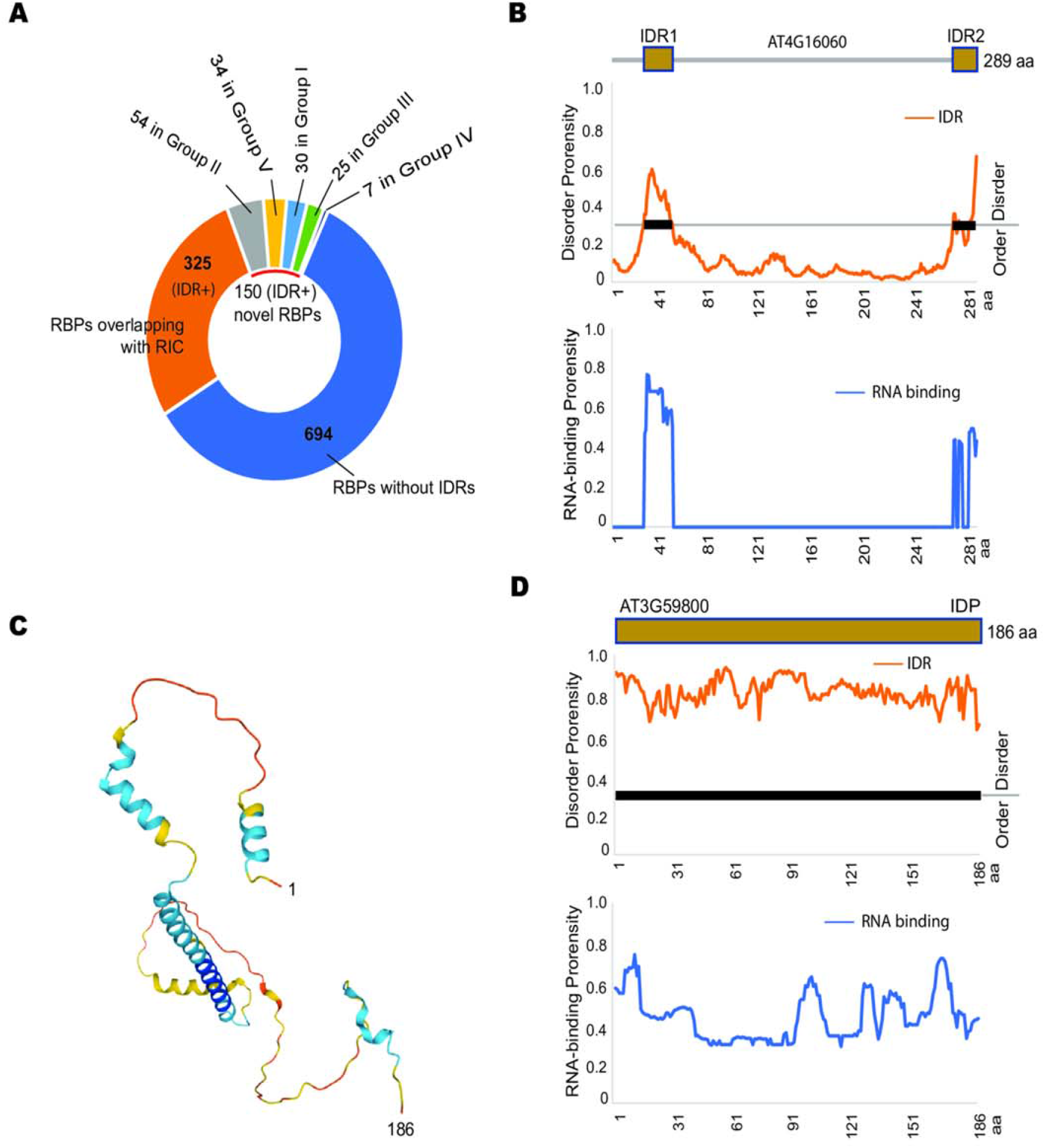
IDRs are engaged in RNA-binding. **(A)** IDRs are distributed in 325 confirmed RBPs and 150 novel RBPs, which were further assigned to 5 groups as indicated in Fig. 4C. The number of RBPs with IDRs are indicated. **(B)** IDRs in RBP (encoded by AT4G16060) may contribute to RNA-binding. Protein domain architecture with two IDRs, the propensity of disorder (orange curve) and RNA-binding (blue curve) for each individual amino acid are shown. Disordered regions are highlighted with thick black lines. **(C)** Predicted structure of RBP, an intrinsically disordered protein (IDP) encoded by AT3G59800. **(D)** Domain architecture, the propensity of disorder (orange curve), and RNA-binding (blue curve) for AT3G59800 are shown.

### Validation of novel RBPs and RBDs

Next, we developed the mRNA pull-down assay to validate the RNA-binding ability of captured novel RBPs/RBDs. To this end, mRNAs (from 12-day-old Arabidopsis leaf tissue) were immobilized to the oligo d(T) magnetic beads as the bait. Recombinant RBPs/RBDs were expressed and purified from *E. coli* and incubated with the bait. The RBP/RBD-RNA interaction was confirmed by western blot. We first tested the efficacy of this method using HLP1, an RRM-type RBP with >5,000 *in vivo* RNA partners (Zhang et al., 2015) as a positive control. As expected, MBP-His-HLP1 was detected only when incubated with beads pre-occupied with mRNA (Figure 6A). Then we applied this method to ATSDH (AT5G51970, Group I) and ESM1 (AT3G14210, Group III). ATSDH is a sorbitol dehydrogenase, and its yeast homolog (SOR1) was classified as RBP by RIC (Matia-Gonzalez et al., 2015). ESM1, which belongs to the GDSL-like lipase/acyl hydrolase superfamily, was one of the four members identified by PPE. It was also discovered by OOPS (Liu et al., 2020). However, neither of these two proteins has been experimentally proven to bind RNA. We showed that beads with mRNAs were able to capture both ATSDH and ESM1, hence validating ATSDH and ESM1 as RBPs (Figure 6B).

**Fig. 6.**
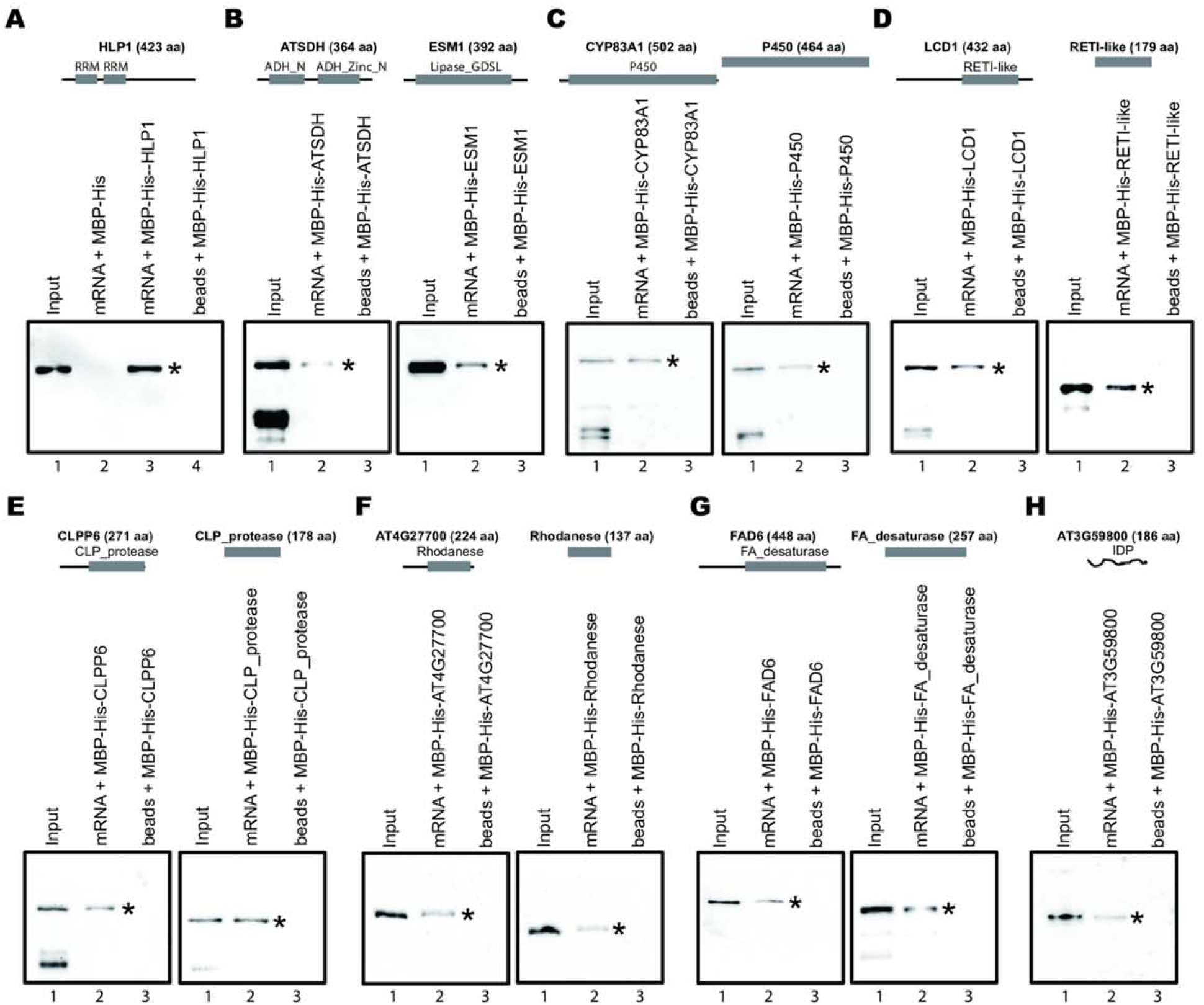
Validation of RNA-binding proteins by an mRNA pull-down assay. **(A)** Establishment of the mRNA pull-down assay by using the known RNA-binding protein HLP1 as the positive control. Recombinant MBP-His-HLP1 was detected by the anti-His antibody only when incubated with mRNA on the oligo d(T)_25_ magnetic beads (lane 3). MBP-His and beads without RNAs were used as negative controls (lanes 2 and 4). (**B)** Full-length ATSDH and ESM1 bind mRNAs *in vitro*. MBP-His-ATSDH/ ESM1 were recognized by the anti-His antibody. (**C)** Full-length CYP83A1 (lane 2, left panel) and its P450 domain (lane 2, right panel) bind to mRNA. (**D)** Full-length LCD1 (left) and its RETICULATA-like (RETI-like) domain interact with mRNA. (**E)** Full-length CLPP6 (left) and its CLP_protease domain interact with mRNA. (**F)** The full-length protein encoded by AT4G27700 (left) and its Rhodanese domain interact with mRNA. (**G)** Full-length FAD6 (left) and its FA_desaturase domain interact with mRNA. (**H)** IDP encoded by AT3G59800 interacts with RNA. All recombinant proteins were indicated with *.

The mRNA pull-down method was further used to validate more RBPs and putative novel RBDs, including three group III RBPs with their active domains (CYP83A1 and its P450 domain shared in the cytochrome P450 family; CLPP6 and its CLP_protease domain shared in the peptidase S14 protein family; LCD1 and its RETICULATA-like domain shared in the RETICULATA protein family), and two Group IV RBPs with their putative RBDs (AT4G27700 and its shared rhodanese-like domain in the rhodanese-like domain-containing protein family; FAD6 and its FA_desaturase domain shared in the fatty acid desaturase family). All the proteins and domains tested here were trapped by mRNA on the beads (Figure 6C-G, lane 2) but not the “beads only” control (Figure 6C-G, lane 3), demonstrating these enzymes as *bona fide* RBPs and suggesting that the active domains of these enzymes have dual roles in both RNA binding and basic metabolic processes. In addition to the moonlighting RBPs, we also confirmed the interaction of AT3G59800, an intrinsically disordered protein (IDP), with RNA (Figure 6H).

Interestingly, we noticed from our raw mass spectrometry data that TGG1 (a myrosinase encoded by AT5G26000) has the most spectral counts in noCL samples across three replicates (Supplemental Data Set 1), therefore was not considered as RBP in our analysis. However, it was considered a candidate RBP in one of the previous RIC studies (Zhang et al., 2016). We showed that TGG1 could bind to oligo d(T) magnetic beads *in vitro* regardless of the presence of mRNA (Supplemental Figure S7, lanes 2 & 3), and such binding could be weakened by stringent washing (Supplemental Figure S7, lane 4), suggesting that TGG1 is more likely to be a non-specific contaminant. In summary, with the mRNA pull-down method, we were able to validate the authenticity of real RBPs and eliminate false-positive interactions.

## Discussion

We have developed the plant phase extraction (PPE) method for isolating RBPs from complex tissues and revealed its effectiveness in identifying the leaf RBPome from Arabidopsis. The unprecedented number of RBPs presented in this study showcased the technical advantages of PPE on multiple levels. For instance, total cell lysate is mixed with Trizol and chloroform throughout phase extraction, which completely destroyed protein-protein interactions to ensure that RBP-RNA adducts are free of non-RBP contaminations and protected RNA from degradation at the same time. Unlike RIC, which relies on oligo d(T) beads affinity purification and extensive washing, PPE does not use beads and thus will not introduce non-specific binding or the bias towards poly(A) RNA-binding proteins. Compared to the previous phase extraction-based method (OOPS) (Liu et al., 2020), PPE captured far more reliable RBPs (1,169 *vs*. 468) and displayed a higher overlap with the 2,701 RBPs combined from all of the RIC studies (58% *vs*. 53%, Figure 4A). When compared with individual RIC studies (Marondedze et al., 2016; Bach-Pages et al., 2020), PPE also generated a more comprehensive leaf RBPome (1,169 *vs*. 236 *vs*. 717, Supplemental Figure S8A), underpinning the success of PPE in isolating RBPs from complex tissue samples. It is worth noting that only 34 RBPs are shared among the three leaf RBPomes, due to the poor overlap of the 4-week-old leaf RBPome with the other two (Supplemental Figure S8A). Notably, our PPE data exhibits a decent overlap with RBPs discovered by the improved RIC (ptRIC), as ∼30% of the RBPs identified by PPE were also captured by ptRIC, accounting for 49% of the RBPs in the 5∼6-week-old leaf RBPome (Bach-Pages et al., 2020). Among the 1,169 PPE-RBPs, 800 were neither in the 4-week-old leaf RBPome, nor in the 5∼6-week-old leaf RBPome (Supplemental Figure S8A). However, they could not be simply ascribed to tissue-specific or developmental-stage-specific RBPs since 278 of them appeared in RBPomes from various cell/tissues types at different development stages and even under stress conditions, such as the 4-day-old etiolated seedling RBPome, the 4∼5-week-old mesophyll protoplast RBPome and the root cell drought-responsive RBPome (Supplemental Figure S8B) (Reichel et al., 2016; Zhang et al., 2016; Marondedze et al., 2019b). Nonetheless, we cannot rule out the possibility that some of the remaining 522 RBPs could be developmental-stage-specific, and others might be uncovered simply because PPE allowed a deep and unbiased identification of RBPs irrespective of RNA types.

In addition to comparing the absolute number of total RBPs in each leaf RBPome, we also focused on the recovery of RBPs associated with rRNA and tRNA to demonstrate the effectiveness of uncovering non-poly(A) RBPs. Altogether, we have identified 160 ribosomal proteins in the PPE-leaf RBPome. Besides a few ribosomal proteins that directly interact with mRNA during translation, many of the ribosomal proteins are structural constituents of the ribosome and bind to rRNA or have extraribosomal functions in RNA metabolism beyond their traditional roles as a structural component and in translation (Warner and McIntosh, 2009; Xiong et al., 2021). Remarkably, PPE identified 41 tRNA ligases/synthetases/methyltransferases involved in translation and tRNA methylation. This is in striking contrast to the other two leaf RBPomes, which only showed seven and five tRNA-related RBPs (Marondedze et al., 2016; Bach-Pages et al., 2020). However, some ribosomal and tRNA-related RBPs in the RIC-leaf RBPome may have been contaminants from non-specific binding of rRNA and tRNA to oligo d(T) beads. The fact that PPE has recovered far more ribosomal and tRNA-related RBPs than any other individual RIC-RBPome further manifested its advantage in capturing non-poly(A) RNA-binding proteins.

In the PPE-leaf RBPome, 51% of the RBPs have no previously assigned roles in RNA biology, which is much higher compared to 25% in RIC-RBPomes (Reichel et al., 2016; Bach-Pages et al., 2020). Furthermore, 44% of the RBPs did not harbor any known RBDs, suggesting that PPE also identified more non-classical RBPs. GO-enrichment analyses showed that these RBPs predominantly participate in numerous metabolic processes, corroborating the notion that various metabolic enzymes moonlight as RBPs (Castello et al., 2015). In some cases, moonlighting enzymes regulate the expression levels of their target transcripts. The crosstalk between gene expression and cellular metabolism mediated by moonlighting enzymes forms the basis of the RNA-enzyme-metabolite (REM) hypothesis (Hentze and Preiss, 2010). A well-known example is glyceraldehyde 3-phosphate dehydrogenase (GAPDH), with dual roles in metabolism and RNA-binding activity. It interacts with tRNA, rRNA, and mRNA to regulate tRNA translocation, mRNA stability, and translation (White and Garcin, 2016). The structure of GAPDH has been well defined, and a large part of the protein is believed to be responsible for RNA-binding. Among the 496 novel RBPs, various metabolic enzymes, such as members of the cytochrome P450 family, peptidase S14 family, rhodanese-like domain-containing protein family, and the fatty acid desaturase family, have no recognizable RBDs. However, many members share conserved motifs in each family, making them excellent candidates for novel RBDs. For example, domain alignment of the fatty acid (FA) desaturases across species displayed FA_desaturase as the only module conserved in all seven FA desaturases, indicating that the FA_desaturase domain may interact directly with RNA (Supplemental Figure S9).

We assigned the 496 novel RBPs into groups based on: if they have RBP homologs in other organisms (Group I); if they have existing RBDs (Group II) or have family members classified as RBP (Group III); if they have other family members co-identified but have not been registered as RBP (Group IV); if proteins do not fit the above-described criteria (Group V). We further developed an mRNA pull-down assay, validated some of the moonlighting enzymes as valid RBPs, and, most importantly, demonstrated a second role of the enzymatic domains as an RBD (Figure 6). This intriguing finding implicates a potential competition or cooperation between RNA and the enzyme-substrate, as RNA binding to the enzymatic domain could either block its access to the substrate or facilitate substrate recruitment. Future work depicting the means of RNA-binding, the RNA partners, and the biological function will shed light on the actual mechanism of these moonlighting enzymes in the intermediary metabolic process, RNA metabolism, and gene regulation.

For the 141 putative RBPs in groups IV and V, we showed that 37 of them contain IDRs that are predicted to be able to interact with RNA (Figure 5 and Supplemental Table 2). IDRs are polypeptide segments lacking hydrophobic amino acids but rich in polar or charged amino acids. In contrast to the classical “structure-function” paradigm that the 3D structure determines the protein function, IDRs are not structural but functional (Dyson and Wright, 2005). A previous study reported that IDRs appear in over 30% of proteins in eukaryotes (Peng et al., 2015). Nearly half of the RNA-binding sites in humans occurred in IDRs (Castello et al., 2016), reflecting IDRs as the hotspots for direct protein-RNA interactions. We also confirmed the direct interaction of an intrinsically disordered protein with RNA by pull-down assay, showing that IDRs contribute to RNA binding.

In summary, we showed that PPE is an efficient and powerful tool to comprehensively identify RBPs from plant tissues. More importantly, it will enable researchers to characterize RBPs and their associated RNAs and pave the way to investigate novel RBP functions under physiological and stress conditions.

## Methods

### Plant growth

Wild-type *Arabidopsis thaliana* in the Columbia-0 background was used. Seeds were surface sterilized with 50% bleach (Clorox) for 5 minutes and rinsed four times in sterilized water. After being stratified at 4 °C for three days, seeds were sown on Murashige and Skoog (MS, pH 5.8) plates containing 1% (v/w) sucrose and 0.8% (w/v) agar and grown at 23 °C under long-day conditions for 12 days. The plates were placed vertically in the chamber to facilitate the collection of roots and leaves separately.

### UV-crosslinking of Arabidopsis tissue

Leaves and roots were separated directly on the plate with a clean blade and transferred to prechilled liquid MS medium (pH 5.8) in 20 mm Petri dishes. The Petri dishes were placed on ice for the entire process. Leaves were crosslinked three times (with a 1-minute interval) at 200 mJ/cm^2^ with 254 nm wavelength in a Hoefer UVC500 Crosslinker (Zhang et al., 2015), with two times of irradiation on the adaxial side and once on the abaxial side as described previously (Bach-Pages et al., 2020). Non-crosslinking control samples (noCL) were soaked in ice-cold liquid MS for approximately the same time as the crosslinked samples (CL). After crosslinking, both noCL and CL tissues were rinsed three times in ice-cold 20 mM Tris-Cl buffer (pH 7.5) and quickly dried with a paper towel. Samples were either immediately frozen in liquid nitrogen and stored at -80 °C or processed directly.

### Plant phase extraction in Arabidopsis leaf tissue

The XRNAX method used in human cell lines (Trendel et al., 2019) was adapted for Arabidopsis tissue with major modifications. Leaf tissues were ground into a fine powder with liquid nitrogen. About 2 g of powder was lysed in 4 ml of lysis buffer (20 mM Tris-Cl, pH 7.5, 0.5 M LiCl, 0.5% LiDS, 0.4% IGEPAL CA630, 2.5% polyvinylpyrrolidone 40, 5 mM DTT, 10 mM Ribonucleoside Vanadyl Complex, 1.5×Roche EDTA-free protease inhibitor) and homogenized by rotating at 4 °C for 30 minutes. Tissue debris was spun down at 10,000 rpm for 15 minutes at 4 °C. The supernatant was transferred to a new tube, centrifuged again at 10,000 rpm for 15 minutes at 4 °C, and the cleared supernatant was transferred to a 50 ml tube. About 400 μl of lysate was saved as input. The rest (∼4 ml) was mixed with 20 ml of Trizol, vigorously vortexed, and incubated at room temperature (RT) for 5 minutes. Then 4 ml of chloroform was added, the mixture was vigorously vortexed and incubated at RT for another 5 minutes, followed by centrifugation at 10,000 g for 15 minutes at 4 °C. After the first round of phase separation, the aqueous phase was removed, and the interphase was transferred to a 1.5 ml tube and centrifuged at 12,000 rpm for 5 minutes at 4 °C. The residual aqueous and organic phases were removed using a syringe with a narrow needle (30G). Then the interphase was subjected to two extra rounds of phase separation, with 1 ml of Trizol and 200 μl of chloroform for each round.

The resulting interphase was washed twice with 1 ml of ethanol and quickly rinsed twice with 1 ml of 50 mM Tris-Cl (pH 7.5). Pellet was spun down at 12,000 rpm for 5 minutes at RT after each wash/rinse. After removing the Tris-Cl buffer from the last rinse, the pellet was resuspended with 300 μl of low SDS buffer (50 mM Tris-Cl, pH7.5, 1 mM EDTA, and 0.1% SDS) by pipetting, then the suspension was spun down with 12,000 rpm for 5 minutes at RT. The supernatant was saved as eluate 1. The pellet was disintegrated again with 300 μl of low SDS buffer and twice with 300 μl of high SDS buffer (50 mM Tris-Cl, pH7.5, 1 mM EDTA, and 0.5% SDS). All four eluates were pooled, and 2 μl of glycogen, NaCl (to a final concentration of 0.3 M), and nine volumes of ethanol were added and mixed well, followed by incubating the mixure at -80 °C for 2 hours. The mix was spun down at maximum speed for 30 minutes at 4 °C and the pellete was rinsed twice with 1 ml of 75% ethanol. The pellet was resuspended with 200 μl of RNase-free water and let sit on ice for 1 hour with occasional pipetting. Then 50 μl of DNase I mixture (25 μl of 10× DNase I buffer, 18 μl of NEB DNase I, 2 μl of 1 M DTT, and 5 μl of RiboLock RNase inhibitor) was added and the solution was incubated at 37 °C for 30 minutes. DNase and other residual protein contaminations (non-RBPs) were further removed by two additional cycles of phase separation as described in the preceding method with 1.25 ml of Trizol and 250 μl of chloroform for each cycle. The final interphase (free of aqueous and organic phases after wash) can be sequentially disintegrated in low SDS buffer, high SDS buffer, precipitated and resuspended again in 100 μl of RNase-free water for downstream applications, or directly resuspended in protein sample buffer for silver staining, Western blot, and mass spectrometry analyses.

### SDS-PAGE, silver staining, and Western blot

Protein samples were mixed well with 4× NuPAGE LDS sample buffer and 10× NuPAGE reducing reagent (both to a final concentration of 1×) and denatured at 99 °C for 10 minutes, before being loaded on a 10% precast NuPAGE Novex Bis-Tirs gel (Thermo Fisher Scientific, Waltham, MA). After electrophoresis at 150 V for an hour and a half, the gel was either rinsed with ultrapure water, fixed and silver-stained according to the manufacturer’s instructions (Pierce™ Silver Stain for Mass Spectrometry, Thermo Fisher Scientific, Waltham, MA) or soaked with nitrocellulose membrane in transfer buffer (25 mM Tris, 190 mM glycine, 20 methanol) before protein transfer. Primary antibodies, anti-AGO1, and anti-histone H3 were purchased from Agrisera; and anti-HLP1 (Zhang et al., 2015) and anti-Fib1/2 were kindly provided by Dr. Xiaofeng Cao at the Institute of Genetics and Developmental Biology, Chinese Academy of Sciences.

### Mass-spectrometry for the PPE-RBPome

#### In-gel digestion

Final interphases were separated using a 4-12% precast NuPAGE Novex Bis-Tris gel. Once the samples ran into the gel (electrophoresis at 150 V for 15 minutes), electrophoresis was stopped, and the gel was stained with Coomassie brilliant blue, then destained and rinsed in double-distilled water. Each gel band was cut with a clean blade and subjected to reduction with 10 mM DTT for 30 min at 60 °C, followed by alkylation with 20 mM iodoacetamide for 45 minutes at room temperature in the dark. Digestion with trypsin was performed at 37 °C overnight (sequencing grade, Cat#90058, Thermo Fisher Scientific, Waltham, MA). Peptides were extracted twice with 5% formic acid, 60% acetonitrile, and vacuum dried.

#### Liquid chromatography-tandem mass spectrometry (LC-MS/MS)

Samples were analyzed by LC-MS using Nano LC-MS/MS (Dionex Ultimate 3000 RLSCnano System, Thermo Fisher Scientific, Waltham, MA) interfaced with Eclipse (Thermo Fisher Scientific, Waltham, MA). Three out of 12.5 μl of the in-gel digested sample were loaded onto a fused silica trap column (Acclaim PepMap 100, 75 μmx2cm, Thermo Fisher Scientific, Waltham, MA). After washing for 5 minutes at 5 μl/min with 0.1% TFA, the trap column was brought in line with an analytical column (Nanoease MZ peptide BEH C18, 130A, 1.7 μm, 75 μm x 250mm, Waters, Milford, MA) for LC-MS/MS. Peptides were fractionated at 300 nL/min using a segmented linear gradient 4-15% B in 30 minutes (where A: 0.2% formic acid, and B: 0.16% formic acid, 80% acetonitrile), 15-25%B in 40 minutes, 25-50% B in 44 minutes, and 50-90% B in 11minutes. Solution B then returns at 4% for 5 minutes for the next run.

The scan sequence began with an MS1 spectrum (Orbitrap analysis, resolution 120,000, scan range from M/Z 375–1500, automatic gain control (AGC) target 1E6, maximum injection time 100 ms). The top S (3 seconds) duty cycle scheme was used for determining the number of MS/MS performed for each cycle. Parent ions of charge 2-7 were selected for MS/MS, and dynamic exclusion of 60 seconds was used to avoid repeat sampling. Parent masses were isolated in the quadrupole with an isolation window of 1.2 m/z, automatic gain control (AGC) target 1E5, and fragmented with higher-energy collisional dissociation with a normalized collision energy of 30%. The fragments were scanned in Orbitrap with a resolution of 15,000. The MS/MS scan ranges were determined by the charge state of the parent ion, but the lower limit was set at 110 atomic mass unit (amu).

#### Database search

The peak list of the LC-MS/MS was generated by Thermo Proteome Discoverer (v. 2.4) into MASCOT Generic Format (MGF) and searched against Araprot 11 and a common lab contaminants (CRAP) database using in house version of X!Tandem (GPM Fury) (Craig and Beavis, 2004). Search parameters are as follows: fragment mass error - 20 ppm; parent mass error - +/- 7 ppm; fixed modification - no fixed modification; variable modifications - methionine monooxidation for the primary search, asparagine deamination, tryptophan oxidation and dioxidation, methionine dioxidation, and glutamine to pyro-glutamine were considered at the refinement stage. Protease specificity: trypsin (C-terminal of Arginine (R)/ Lysine (K) unless followed by Proline (P)) with one missed cleavage during the preliminary search and five missed cleavages during refinement. Minimum acceptable peptide and protein expectation scores were set at 10^−2^ and 10^−4^, respectively. The overall peptide false-positive rate was 0.07% (Gupta et al., 2011).

### Data analysis

#### Definition of RBPs identified by PPE

Proteins with CL > 1 and fold change [CL/noCL] ≥ 2 in at least two out of three replicates were considered in statistical analysis. Statistical analysis was performed as described with modifications (Backlund et al., 2020). Data of qualified protein were first cleaned using the limma package to remove batch effects (Ritchie et al., 2015). The CL and noCL data were then normalized separately with a variance stabilization normalization (vsn) approach using the vsn package (Huber et al., 2002). Next, differential expression analysis was carried out by the limma package. Proteins with a fold change [CL/noCL] ≥ 2 at an adjusted *p*-value (FDR) < 5% were defined as RBPs, and those with a fold change [CL/noCL] ≥ 1.5 at an FDR < 20% were considered as candidate RBPs.

#### GO enrichment analysis

The enrichment of molecular functions, biological processes, and cellular components was analyzed using Metascape (Zhou et al., 2019). Terms with *p*-value <0.01were considered enriched.

#### Pfam domain annotation and enrichment analysis

One representative isoform of each gene was retrieved from the Arabidopsis genome (Araport 11) on the TAIR website (https://www.arabidopsis.org/). Protein sequence search for Pfam-A domain was conducted with the domain-search algorithm HMMER3, using an E-value < 1.0 as the cut-off. Over-representation of a Pfam domain was tested with Fisher’s exact test, and *p*-values were corrected using the Benjamini and Hochberg method.

#### IDR and function prediction

The predicted Arabidopsis IDR dataset was downloaded from MobiDB (Piovesan et al., 2021). Gene IDs of the PPE-RBPome were converted into Uniprot IDs and searched against the IDR dataset to retrieve IDRs residing in the PPE-RBPome. RNA-IDR interaction was predicted using flDPnn (Hu et al., 2021).

### Validation of RBPs

#### DNA constructs, protein expression, and purification

Full-length CDS of candidate RBPs and DNA fragments covering the RBDs were PCR amplified from cDNAs reverse-transcribed from Col-0 mRNA using the Phusion DNA Polymerase (NEB), and cloned into the pMCSG9 vector using the ligation-independent cloning (LIC) strategy as described (Eschenfeldt et al., 2009). Plasmids containing the *His-MBP-RBP/RBD* fusion constructs were transformed into *E. coli* BL21 (RIL) strain for expression. Proteins were purified using the Ni-NTA His-Bind Resin following the manufacturer’s instructions (MilliporeSigma, Burlington, MA). See Supplemental Data Set 8 for a list of primers.

#### mRNA pull-down

For each pull-down assay, 25 μg of total RNAs from 12-day-old Arabidopsis leaves were extracted using the Trizol method and incubated with 30 μl of pre-equilibrated oligo d(T)_25_ magnetic beads (NEB) for 15 minutes at room temperature with continuous rotation. Then the beads were briefly washed once in lysis/binding buffer (100 mM Tris-Cl, pH7.5, 0.5 M LiCl, 0.5% LiDS, 1 mM EDTA, 5 mM DTT), wash buffer I (20 mM Tris-Cl, pH7.5, 0.5 M LiCl, 0.1% LiDS, 1 mM EDTA, 5 mM DTT), wash buffer II (20 mM Tris-Cl, pH7.5, 0.5 M LiCl, 1 mM EDTA, 5 mM DTT), respectively, and three times in pull-down binding buffer (20 mM Tris-Cl, pH7.5, 150 mM NaCl, 5 mM MgCl_2_, 0.1% IGEPAL CA-630 and 5 mM DTT). After the final wash, 200 μg of BSA diluted in 400 μl of pull-down binding buffer was added to the beads and incubated at 4 °C for 1 hour with continuous rotation. Then 3 μg of purified recombinant protein was added with 4 μl of RiboLock RNase Inhibitor (Thermo Fisher Scientific, Waltham, MA). The mixture was incubated at room temperature for 1 hour, followed by washing in pull-down wash buffer (20 mM Tris-Cl, pH7.5, 150 mM NaCl, 0.2% IGEPAL CA-630) for four times, with a brief vortex in each wash. The final beads were added with 30 μl of 1 x SDS protein loading buffer and incubated at 99 °C for 10 minutes with vigorous shaking. Ten μl of the final elution were used for SDS-PAGE gel separation and Western blot analysis. Candidate RBPs were detected with anti-His monoclonal antibody (MilliporeSigma, Burlington, MA).

## Acknowledgments

**This research was funded by the United States Department of Agriculture-Agricultural Research Service, National Program 301: Plant Genetic Resources, Genomics, and Genetic Improvement (project number 2036-13210-012-00-D) and the National Institute of Food and Agriculture (project number CA-R-BPS-5084-H). The authors thank Dr. Xiaofeng Cao for providing anti-HLP1 and anti-Fib1/2 antibodies and Dr. Haiyan Zheng for mass-spectrometry analyses (Center for Advanced Biotechnology and Medicine, Rutgers University)**.

## Author Contributions

**Y.Z. and D.S. designed the project. D.S. and X.C. supervised the experiments. Y.Z. performed the experiments and data analysis**.

**Y.X carried out the statistical analysis of the mass-spectrometry data and retrieved and created the Arabidopsis RBD reference database. T.S**., **J.F**., **and X.C. provided conceptual advice and assistance with experiments. Y.Z. and D.S. wrote the manuscript with joint contributions from all the authors. All authors have read and approved the manuscript**.

## Parsed Citations

Albaqami, M., Laluk, K., and Reddy, A.S.N. (2019). The Arabidopsis splicing regulator SR45 confers salt tolerance in a splice isoform-dependent manner. Plant Mol Biol 100, 379–390.

Bach-Pages, M., Homma, F., Kourelis, J., Kaschani, F., Mohammed, S., Kaiser, M., van der Hoorn, R.A.L., Castello, A., and Preston, G.M. (2020). Discovering the RNA-Binding Proteome of Plant Leaves with an Improved RNAInteractome Capture Method. Biomolecules 10.

Backlund, M., Stein, F., Rettel, M., Schwarzl, T., Perez-Perri, J.I., Brosig, A., Zhou, Y., Neu-Yilik, G., Hentze, M.W., and Kulozik, A.E. (2020). Plasticity of nuclear and cytoplasmic stress responses of RNA-binding proteins. Nucleic acids research 48, 4725–4740.

Baltz, A.G., Munschauer, M., Schwanhausser, B., Vasile, A., Murakawa, Y., Schueler, M., Youngs, N., Penfold-Brown, D., Drew, K., Milek, M., Wyler, E., Bonneau, R., Selbach, M., Dieterich, C., and Landthaler, M. (2012). The mRNA-bound proteome and its global occupancy profile on protein-coding transcripts. Mol Cell 46, 674–690.

Bao, X., Guo, X., Yin, M., Tariq, M., Lai, Y., Kanwal, S., Zhou, J., Li, N., Lv, Y., Pulido-Quetglas, C., Wang, X., Ji, L., Khan, M.J., Zhu, X., Luo, Z., Shao, C., Lim, D.H., Liu, X., Li, N., Wang, W., He, M., Liu, Y.L., Ward, C., Wang, T., Zhang, G., Wang, D., Yang, J., Chen, Y., Zhang, C., Jauch, R., Yang, Y.G., Wang, Y., Qin, B., Anko, M.L., Hutchins, A.P., Sun, H., Wang, H., Fu, X.D., Zhang, B., and Esteban, M.A. (2018). Capturing the interactome of newly transcribed RNA. Nat Methods 15, 213–220.

Bechara, E.G., Sebestyen, E., Bernardis, I., Eyras, E., and Valcarcel, J. (2013). RBM5, 6, and 10 differentially regulate NUMB alternative splicing to control cancer cell proliferation. Mol Cell 52, 720–733.

Beckmann, B.M., Horos, R., Fischer, B., Castello, A., Eichelbaum, K., Alleaume, A.M., Schwarzl, T., Curk, T., Foehr, S., Huber, W., Krijgsveld, J., and Hentze, M.W. (2015). The RNA-binding proteomes from yeast to man harbour conserved enigmRBPs. Nat Commun 6, 10127.

Bush, M.S., Crowe, N., Zheng, T., and Doonan, J.H. (2015). The RNAhelicase, eIF4A-1, is required for ovule development and cell size homeostasis in Arabidopsis. Plant J 84, 989–1004.

Calabretta, S., and Richard, S. (2015). Emerging Roles of Disordered Sequences in RNA-Binding Proteins. Trends Biochem Sci 40, 662–672.

Castello, A., Hentze, M.W., and Preiss, T. (2015). Metabolic Enzymes Enjoying New Partnerships as RNA-Binding Proteins. Trends Endocrinol Metab 26, 746–757.

Castello, A., Fischer, B., Frese, C.K., Horos, R., Alleaume, A.M., Foehr, S., Curk, T., Krijgsveld, J., and Hentze, M.W. (2016). Comprehensive Identification of RNA-Binding Domains in Human Cells. Mol Cell 63, 696–710.

Castello, A., Fischer, B., Eichelbaum, K., Horos, R., Beckmann, B.M., Strein, C., Davey, N.E., Humphreys, D.T., Preiss, T., Steinmetz, L.M., Krijgsveld, J., and Hentze, M.W. (2012). Insights into RNAbiology from an atlas of mammalian mRNA-binding proteins. Cell 149, 1393–1406.

Chang, C.H., Curtis, J.D., Maggi, L.B., Jr., Faubert, B., Villarino, A.V., O’Sullivan, D., Huang, S.C., van der Windt, G.J., Blagih, J., Qiu, J., Weber, J.D., Pearce, E.J., Jones, R.G., and Pearce, E.L. (2013). Posttranscriptional control of T cell effector function by aerobic glycolysis. Cell 153, 1239–1251.

Cheng, S., Gutmann, B., Zhong, X., Ye, Y., Fisher, M.F., Bai, F., Castleden, I., Song, Y., Song, B., Huang, J., Liu, X., Xu, X., Lim, B.L., Bond, C.S., Yiu, S.M., and Small, I. (2016). Redefining the structural motifs that determine RNAbinding and RNAediting by pentatricopeptide repeat proteins in land plants. Plant J 85, 532–547.

Chu, E., Koeller, D.M., Casey, J.L., Drake, J.C., Chabner, B.A., Elwood, P.C., Zinn, S., and Allegra, C.J. (1991). Autoregulation of human thymidylate synthase messenger RNAtranslation by thymidylate synthase. Proc Natl Acad Sci U S A 88, 8977–8981.

Craig, R., and Beavis, R.C. (2004). TANDEM: matching proteins with tandem mass spectra. Bioinformatics 20, 1466–1467.

Deleris, A., Gallego-Bartolome, J., Bao, J., Kasschau, K.D., Carrington, J.C., and Voinnet, O. (2006). Hierarchical action and inhibition of plant Dicer-like proteins in antiviral defense. Science 313, 68–71.

Despic, V., Dejung, M., Gu, M., Krishnan, J., Zhang, J., Herzel, L., Straube, K., Gerstein, M.B., Butter, F., and Neugebauer, K.M. (2017). Dynamic RNA-protein interactions underlie the zebrafish maternal-to-zygotic transition. Genome Res 27, 1184–1194.

Dyson, H.J., and Wright, P.E. (2005). Intrinsically unstructured proteins and their functions. Nat Rev Mol Cell Biol 6, 197–208.

Eschenfeldt, W.H., Lucy, S., Millard, C.S., Joachimiak, A., and Mark, I.D. (2009). Afamily of LIC vectors for high-throughput cloning and purification of proteins. Methods Mol Biol 498, 105–115.

Gebauer, F., Schwarzl, T., Valcarcel, J., and Hentze, M.W. (2021). RNA-binding proteins in human genetic disease. Nat Rev Genet 22, 185–198.

Gerstberger, S., Hafner, M., and Tuschl, T. (2014). Acensus of human RNA-binding proteins. Nat Rev Genet 15, 829–845.

Glisovic, T., Bachorik, J.L., Yong, J., and Dreyfuss, G. (2008). RNA-binding proteins and post-transcriptional gene regulation. FEBS Lett 582, 1977–1986.

Gupta, N., Bandeira, N., Keich, U., and Pevzner, P.A. (2011). Target-decoy approach and false discovery rate: when things may go wrong. J Am Soc Mass Spectrom 22, 1111–1120.

Heintzen, C., Nater, M., Apel, K., and Staiger, D. (1997). AtGRP7, a nuclear RNA-binding protein as a component of a circadian-regulated negative feedback loop in Arabidopsis thaliana. Proc Natl Acad Sci U S A 94, 8515–8520.

Hentze, M.W., and Argos, P. (1991). Homology between IRE-BP, a regulatory RNA-binding protein, aconitase, and isopropylmalate isomerase. Nucleic acids research 19, 1739–1740.

Hentze, M.W., and Preiss, T. (2010). The REM phase of gene regulation. Trends Biochem Sci 35, 423–426.

Hentze, M.W., Castello, A., Schwarzl, T., and Preiss, T. (2018). Abrave new world of RNA-binding proteins. Nat Rev Mol Cell Biol 19, 327–341.

Hu, G., Katuwawala, A., Wang, K., Wu, Z., Ghadermarzi, S., Gao, J., and Kurgan, L. (2021). flDPnn: Accurate intrinsic disorder prediction with putative propensities of disorder functions. Nat Commun 12, 4438.

Huang, R., Han, M., Meng, L., and Chen, X. (2018). Transcriptome-wide discovery of coding and noncoding RNA-binding proteins. Proc Natl Acad Sci U S A 115, E3879–E3887.

Huber, W., von Heydebreck, A., Sultmann, H., Poustka, A., and Vingron, M. (2002). Variance stabilization applied to microarray data calibration and to the quantification of differential expression. Bioinformatics 18 Suppl 1, S96–104.

Kim, Y.O., Kim, J.S., and Kang, H. (2005). Cold-inducible zinc finger-containing glycine-rich RNA-binding protein contributes to the enhancement of freezing tolerance in Arabidopsis thaliana. Plant J 42, 890–900.

Kwon, S.C., Yi, H., Eichelbaum, K., Fohr, S., Fischer, B., You, K.T., Castello, A., Krijgsveld, J., Hentze, M.W., and Kim, V.N. (2013). The RNA-binding protein repertoire of embryonic stem cells. Nat Struct Mol Biol 20, 1122–1130.

Lefebvre, S., Burglen, L., Reboullet, S., Clermont, O., Burlet, P., Viollet, L., Benichou, B., Cruaud, C., Millasseau, P., Zeviani, M., and et al. (1995). Identification and characterization of a spinal muscular atrophy-determining gene. Cell 80, 155–165.

Liao, Y., Castello, A., Fischer, B., Leicht, S., Foehr, S., Frese, C.K., Ragan, C., Kurscheid, S., Pagler, E., Yang, H., Krijgsveld, J., Hentze, M.W., and Preiss, T. (2016). The Cardiomyocyte RNA-Binding Proteome: Links to Intermediary Metabolism and Heart Disease. Cell Rep 16, 1456–1469.

Liepelt, A., Naarmann-de Vries, I.S., Simons, N., Eichelbaum, K., Fohr, S., Archer, S.K., Castello, A., Usadel, B., Krijgsveld, J., Preiss, T., Marx, G., Hentze, M.W., Ostareck, D.H., and Ostareck-Lederer, A. (2016). Identification of RNA-binding Proteins in Macrophages by Interactome Capture. Mol Cell Proteomics 15, 2699–2714.

Liu, J., Zhang, C., Jia, X., Wang, W., and Yin, H. (2020). Comparative analysis of RNA-binding proteomes under Arabidopsis thaliana-Pst DC3000-PAMP interaction by orthogonal organic phase separation. Int J Biol Macromol 160, 47–54.

Lorkovic, Z.J., and Barta, A. (2002). Genome analysis: RNArecognition motif (RRM) and K homology (KH) domain RNA-binding proteins from the flowering plant Arabidopsis thaliana. Nucleic acids research 30, 623–635.

Louis, J.M., Agarwal, A., Aduri, R., and Talukdar, I. (2021). Global analysis of RNA-protein interactions in TNF-alpha induced alternative splicing in metabolic disorders. FEBS Lett 595, 476–490.

Lunde, B.M., Moore, C., and Varani, G. (2007). RNA-binding proteins: modular design for efficient function. Nat Rev Mol Cell Biol 8, 479–490.

Macknight, R., Bancroft, I., Page, T., Lister, C., Schmidt, R., Love, K., Westphal, L., Murphy, G., Sherson, S., Cobbett, C., and Dean, C. (1997). FCA, a gene controlling flowering time in Arabidopsis, encodes a protein containing RNA-binding domains. Cell 89, 737–745.

Marondedze, C. (2020). The increasing diversity and complexity of the RNA-binding protein repertoire in plants. Proc Biol Sci 287, 20201397.

Marondedze, C., Thomas, L., Lilley, K.S., and Gehring, C. (2019a). Drought Stress Causes Specific Changes to the Spliceosome and Stress Granule Components. Front Mol Biosci 6, 163.

Marondedze, C., Thomas, L., Gehring, C., and Lilley, K.S. (2019b). Changes in the Arabidopsis RNA-binding proteome reveal novel stress response mechanisms. BMC Plant Biol 19, 139.

Marondedze, C., Thomas, L., Serrano, N.L., Lilley, K.S., and Gehring, C. (2016). The RNA-binding protein repertoire of Arabidopsis thaliana. Sci Rep 6, 29766.

Matia-Gonzalez, A.M., Laing, E.E., and Gerber, A.P. (2015). Conserved mRNA-binding proteomes in eukaryotic organisms. Nat Struct Mol Biol 22, 1027–1033.

Patel, A., Lee, H.O., Jawerth, L., Maharana, S., Jahnel, M., Hein, M.Y., Stoynov, S., Mahamid, J., Saha, S., Franzmann, T.M., Pozniakovski, A., Poser, I., Maghelli, N., Royer, L.A., Weigert, M., Myers, E.W., Grill, S., Drechsel, D., Hyman, A.A., and Alberti, S. (2015). ALiquid-to-Solid Phase Transition of the ALS Protein FUS Accelerated by Disease Mutation. Cell 162, 1066–1077.

Peng, Z., Yan, J., Fan, X., Mizianty, M.J., Xue, B., Wang, K., Hu, G., Uversky, V.N., and Kurgan, L. (2015). Exceptionally abundant exceptions: comprehensive characterization of intrinsic disorder in all domains of life. Cell Mol Life Sci 72, 137–151.

Piovesan, D., Necci, M., Escobedo, N., Monzon, A.M., Hatos, A., Micetic, I., Quaglia, F., Paladin, L., Ramasamy, P., Dosztanyi, Z., Vranken, W.F., Davey, N.E., Parisi, G., Fuxreiter, M., and Tosatto, S.C.E. (2021). MobiDB: intrinsically disordered proteins in 2021. Nucleic acids research 49, D361–D367.

Queiroz, R.M.L., Smith, T., Villanueva, E., Marti-Solano, M., Monti, M., Pizzinga, M., Mirea, D.M., Ramakrishna, M., Harvey, R.F., Dezi, V., Thomas, G.H., Willis, A.E., and Lilley, K.S. (2019). Comprehensive identification of RNA-protein interactions in any organism using orthogonal organic phase separation (OOPS). Nat Biotechnol 37, 169–178.

Rakitina, D.V., Taliansky, M., Brown, J.W., and Kalinina, N.O. (2011). Two RNA-binding sites in plant fibrillarin provide interactions with various RNAsubstrates. Nucleic Acids Res 39, 8869–8880.

Reichel, M., Liao, Y., Rettel, M., Ragan, C., Evers, M., Alleaume, A.M., Horos, R., Hentze, M.W., Preiss, T., and Millar, A.A. (2016). In Planta Determination of the mRNA-Binding Proteome of Arabidopsis Etiolated Seedlings. Plant Cell 28, 2435–2452.

Ritchie, M.E., Phipson, B., Wu, D., Hu, Y., Law, C.W., Shi, W., and Smyth, G.K. (2015). limma powers differential expression analyses for RNA-sequencing and microarray studies. Nucleic acids research 43, e47.

Schomburg, F.M., Patton, D.A., Meinke, D.W., and Amasino, R.M. (2001). FPA, a gene involved in floral induction in Arabidopsis, encodes a protein containing RNA-recognition motifs. Plant Cell 13, 1427–1436.

Silverman, I.M., Li, F., and Gregory, B.D. (2013). Genomic era analyses of RNAsecondary structure and RNA-binding proteins reveal their significance to post-transcriptional regulation in plants. Plant Sci 205-206, 55–62.

Singh, G., Pratt, G., Yeo, G.W., and Moore, M.J. (2015). The clothes make the mRNA: Past and present trends in mRNP Fashion. Annu. Rev. Biochem. 84, 325–354.

Song, J.J., Liu, J., Tolia, N.H., Schneiderman, J., Smith, S.K., Martienssen, R.A., Hannon, G.J., and Joshua-Tor, L. (2003). The crystal structure of the Argonaute2 PAZ domain reveals an RNAbinding motif in RNAi effector complexes. Nat Struct Biol 10, 1026–1032.

Sysoev, V.O., Fischer, B., Frese, C.K., Gupta, I., Krijgsveld, J., Hentze, M.W., Castello, A., and Ephrussi, A. (2016). Global changes of the RNA-bound proteome during the maternal-to-zygotic transition in Drosophila. Nat Commun 7, 12128.

Trendel, J., Schwarzl, T., Horos, R., Prakash, A., Bateman, A., Hentze, M.W., and Krijgsveld, J. (2019). The Human RNA-Binding Proteome and Its Dynamics during Translational Arrest. Cell 176, 391–403 e319.

Urdaneta, E.C., Vieira-Vieira, C.H., Hick, T., Wessels, H.H., Figini, D., Moschall, R., Medenbach, J., Ohler, U., Granneman, S., Selbach, M., and Beckmann, B.M. (2019). Purification of cross-linked RNA-protein complexes by phenol-toluol extraction. Nat Commun 10, 990.

Verkerk, A.J., Pieretti, M., Sutcliffe, J.S., Fu, Y.H., Kuhl, D.P., Pizzuti, A., Reiner, O., Richards, S., Victoria, M.F., Zhang, F.P., and et al. (1991). Identification of a gene (FMR-1) containing a CGG repeat coincident with a breakpoint cluster region exhibiting length variation in fragile X syndrome. Cell 65, 905–914.

Wang, X., Liu, R., Zhu, W., Chu, H., Yu, H., Wei, P., Wu, X., Zhu, H., Gao, H., Liang, J., Li, G., and Yang, W. (2019). UDP-glucose accelerates SNAI1 mRNA decay and impairs lung cancer metastasis. Nature 571, 127–131.

Warner, J.R., and McIntosh, K.B. (2009). How common are extraribosomal functions of ribosomal proteins? Mol Cell 34, 3–11.

White, M.R., and Garcin, E.D. (2016). The sweet side of RNAregulation: glyceraldehyde-3-phosphate dehydrogenase as a noncanonical RNA-binding protein. Wiley Interdiscip Rev RNA 7, 53–70.

Xiong, W., Lan, T., and Mo, B. (2021). Extraribosomal Functions of Cytosolic Ribosomal Proteins in Plants. Front Plant Sci 12, 607157.

Zhang, X., Zhao, H., Gao, S., Wang, W.C., Katiyar-Agarwal, S., Huang, H.D., Raikhel, N., and Jin, H. (2011). Arabidopsis Argonaute 2 regulates innate immunity via miRNA393(*)-mediated silencing of a Golgi-localized SNARE gene, MEMB12. Mol Cell 42, 356–366.

Zhang, Y., Cheng, Y.T., Bi, D., Palma, K., and Li, X. (2005). MOS2, a protein containing G-patch and KOW motifs, is essential for innate immunity in Arabidopsis thaliana. Curr Biol 15, 1936–1942.

Zhang, Y., Gu, L., Hou, Y., Wang, L., Deng, X., Hang, R., Chen, D., Zhang, X., Zhang, Y., Liu, C., and Cao, X. (2015). Integrative genome-wide analysis reveals HLP1, a novel RNA-binding protein, regulates plant flowering by targeting alternative polyadenylation. Cell Res 25, 864–876.

Zhang, Z., Boonen, K., Ferrari, P., Schoofs, L., Janssens, E., van Noort, V., Rolland, F., and Geuten, K. (2016). UV crosslinked mRNA-binding proteins captured from leaf mesophyll protoplasts. Plant Methods 12, 42.

Zhou, Y., Zhou, B., Pache, L., Chang, M., Khodabakhshi, A.H., Tanaseichuk, O., Benner, C., and Chanda, S.K. (2019). Metascape provides a biologist-oriented resource for the analysis of systems-level datasets. Nat Commun 10, 1523.

